# Intermediate Relative Humidity Preserves Respiratory Syncytial Virus via a Semi-Solid Bioaerosol State

**DOI:** 10.64898/2026.05.28.728343

**Authors:** Yuhui Guo, Deepak Sapkota, Ajay Sajan, HoangDinh Huynh, Imtiaz Taimoor, Jeffrey Kahn, Hui Ouyang

## Abstract

Respiratory syncytial virus (RSV) transmission via the aerosol route remains poorly understood, particularly with respect to how evolving virus-laden particles (bioaerosols) microenvironments influence viral survival. Bioaerosol particles contain complex mixtures of organic and inorganic components, and their physicochemical properties change dynamically during evaporation as water is lost upon emission from respiratory activities. These changes directly affect the local environment surrounding embedded virus during both the evaporation stage and the subsequent equilibrium state. However, how these microenvironmental conditions under different relative humidity (RH) levels regulate RSV survival remains unclear.

In this study, we quantified RSV survival during the evaporation and early equilibrium stages using a flow-tube system with controlled residence times. Bioaerosols were generated from virus medium alone or supplemented with bovine serum albumin (BSA) or mucin and evaluated under low (35%) and intermediate (61%) RH conditions. Viral infectivity was normalized to RNA copy number to account for particle and sampling losses. At 35% RH, RSV infectivity decreased by one to three orders of magnitude, depending on the solution composition. In contrast, survival was significantly higher at intermediate RH, particularly for virus medium and BSA-supplemented aerosols. Scanning electron microscopy revealed that low RH conditions promote efflorescence, whereas intermediate RH results in viscous or semi-solid particles with higher water content. These observations suggest that efflorescence is associated with enhanced RSV inactivation, while viscous or semi-solid phases tend to preserve RSV in the aerosol state for respirable particles. Overall, RSV infectivity depends strongly on particle chemical composition, phase state (effloresced versus semi-solid), and relative humidity. These results highlight the importance of characterizing particle phase behavior and chemical composition during early aerosol processes to improve mechanistic understanding of viral survival relevant to short-range transmission.

## 1. Introduction

Airborne virus-laden bioaerosols have been responsible for some of the most devastating, costly, and difficult-to-control respiratory diseases in humans and animals, including respiratory syncytial virus (RSV), SARS-CoV-2, and influenza virus in humans^1^, as well as porcine reproductive and respiratory syndrome (PRRS) virus in animals^2,3^. Among these pathogens, RSV is of particular concern because it causes recurrent seasonal infections and severe lower respiratory tract disease, especially in infants, older adults, and individuals with compromised immune or respiratory function^4^. Although RSV vaccines^5^ and monoclonal antibody–based prophylactic interventions have recently been introduced, their availability does not eliminate the need to understand airborne transmission, as population-level coverage, durability of protection, and impacts on infection and onward transmission continue to evolve.

Droplets generated by respiratory activities undergo rapid evaporation upon emission and subsequently reach equilibrium with the surrounding environment. Viral inactivation rates differ between the evaporation and equilibrium stages^6^, likely due to rapid and dynamic changes in microenvironmental conditions within the particle. Viruses have been reported to decay more rapidly during the evaporation stage^7^. Although evaporation typically occurs over only a few seconds, particles in this transient stage can contribute substantially to the higher-risk short-range (∼2 m) transmission due to both the high velocities associated with speaking, coughing, and sneezing^8,9^ and potential thermal plume generated along with respiratory activities^10^.

Respiratory aerosol particles provide complex organic–inorganic microenvironments that strongly influence viral survival. These particles contain a range of components, including salts, proteins, carbohydrates, surfactants, lipids, and other respiratory-fluid constituents^11–13^. Immediately after aerosol generation, rapid water loss concentrates these nonvolatile components, leading to dynamic changes in microenvironmental conditions, such as particle size, viscosity, phase state, pH, intraparticle chemical distributions, and solute activity surrounding embedded virions ^14,15^. Consistent with these evolving conditions, previous studies have shown that viral persistence depends strongly on the composition of the suspending medium, with notable differences observed between human saliva, artificial saliva, airway extracellular material, and cell culture media^16–18^.

Despite extensive experimental observations of viral survival in bioaerosols, mechanistic understanding remains limited due to insufficient characterization of the evolving microenvironment experienced by viruses within aerosol particles. Existing studies have examined larger droplets in both suspended and deposited states to improve mechanistic insight. In suspended droplets, protein-rich components such as mucin can enhance viral survival by modifying evaporation kinetics, increasing viscosity, limiting molecular diffusion, or promoting semisolid or glassy states that reduce chemical stress on virions^19^. In deposited droplets^20^, colocalization of viruses with proteins has been shown to further enhance survival. Together, these studies provide an important foundation ^21–23^; however, additional studies are needed to better characterize the microenvironmental conditions governing viral survival across different aerosolization solutions, as these effects are strongly dependent on chemical composition, particularly for respirable particles (<5 μm) where aerosol transmission is most relevant.

Moreover, compared with influenza and other respiratory viruses, RSV-laden bioaerosol survival measurements are extremely scarce. Until recently, Niazi et al.^24^ measured RSV survival ratios at three RH conditions and under dynamically changing RH using a rotating drum and reported that RSV survival during early suspension periods (< 5 min) exhibited little dependence on RH. While these findings provide important benchmarks, they do not resolve how RSV survival in the evaporation stage, nor how protein constituents of respiratory fluids modulate RSV stability.

In this study, we quantify RSV survival during the evaporation and early equilibrium stages of the airborne period, aiming to provide guidance for RSV short-range transmission. We developed a flow-tube aerosolization system that enables sampling of RSV-laden bioaerosol particles at three representative time points: immediately after aerosolization (*t* = 0 s), during evaporation (*t* = 10 s), and equilibrium (*t* = 70 s) in early suspension. Bioaerosols were generated from virus medium either alone or supplemented separately with 3 g L^−1^ BSA, or 3 g L^−1^ mucin. RSV survival ratios were measured under two representative indoor relative humidity (RH) conditions (∼35% and ∼61%), corresponding to typical winter and summer environments in temperate regions. In addition, particle phase state and intraparticle chemical distributions were characterized using SEM imaging to provide mechanistic insight into observed RSV survival trends.

### 2.1 Methods and Materials

#### RSV-laden Bioaerosol Generation and Collection

All experiments involving viable RSV were conducted in a biosafety level-2 (BSL-2) laboratory using two custom-built negative-pressure chambers (24 in × 24 in × 48 in) that fully enclosed the aerosolization system. As shown in **Figure 1**, RSV-laden bioaerosols were generated using a custom-built Large Aerosol Generator (LAG) coupled with a syringe pump (Braintree Scientific, Inc., MA, USA). The LAG was positioned inside a stainless-steel flow tube (4 in outer diameter; McMaster sanitary fittings; total length is 90 in). Four inlet ports introduced dilution air into the system, with or without pre-conditioned relative humidity (RH) control, resulting in a total volumetric flow rate of 25.5 ± 0.3 L min^−1^, as measured by a flow meter (Model 4040, TSI, MN, USA).

**Figure 1.**
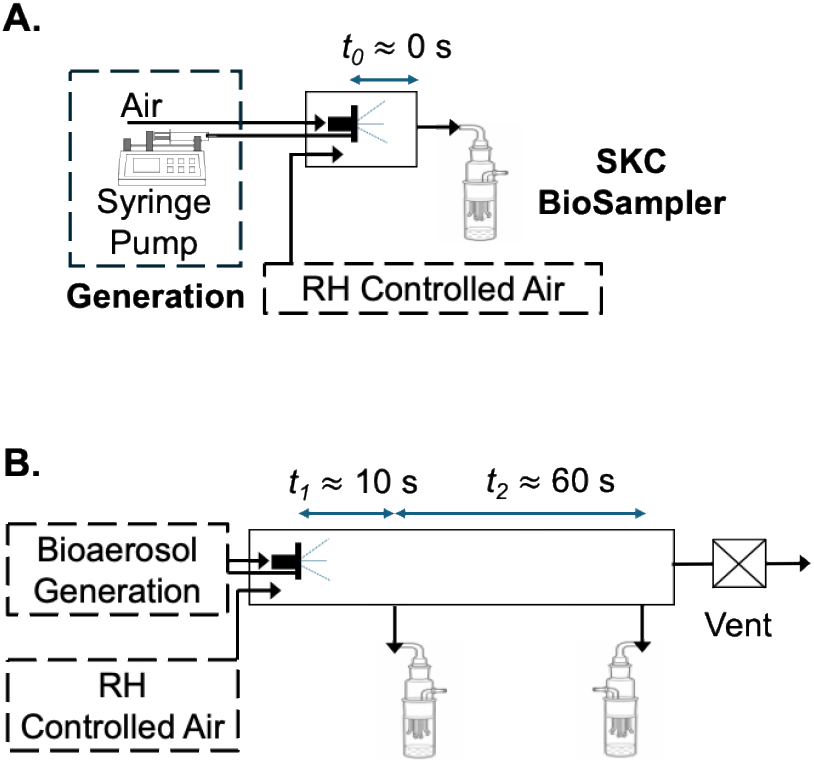
Experimental setup for measuring RSV survival in bioaerosols during (A) aerosolization and (B) evaporation and equilibrium stages, where ***t***_**1**_ ≈ 10s and ***t***_**2**_ ≈ 60s.

The syringe pump delivered the suspension medium to the LAG nozzle at a flow rate of 0.18– 0.25 mL min^−1^. The syringe flow rate was adjusted to achieve the desired RH within the flow tube, while the total aerosolization volume was held constant at 2.5 mL for all experiments. Temperature and RH were continuously monitored using probes (Ahlborn, Germany, Model: FHAD46-Cx) positioned at both upstream and downstream sampling locations. Aerosol generation was initiated and allowed to proceed until RH stabilized at both locations prior to sampling. The difference in RH between the two sampling locations was consistently within ∼4%.

BioSamplers (SKC Inc., Part No. 225-9593) were operated under standard conditions at a sampling flow rate of 12.5 L min^−1^ and collected bioaerosols for 10 min. A liquid trap was installed downstream of each BioSampler to prevent collection medium from being drawn into the filter. Viral transport medium (Hardy Diagnostic, 460029P) was used as the collection medium with an initial volume of 5 mL, and the mass of the VTM was measured before and after sampling to make sure samplings were consistent.

#### Aerosolization, Evaporation and Equilibrium

Two configurations were employed to separate the aerosolization stage from the evaporation and equilibrium stages. In the first configuration (Fig. 1A), the sampling port of the SKC BioSampler was positioned close to the LAG outlet (approximately 2 cm away) to enable direct sampling of freshly aerosolized droplets with a negligible residence time. The relative humidity at this point was monitored at ∼91%. In the second configuration (Fig. 1B), two sampling ports were positioned to provide average residence times of approximately 10 s and 70 s, respectively, corresponding to a total flow rate of 25.5 LPM and a travel distance of 24 in and a flow rate of 13 LPM and a travel distance of 66 in.

Accordingly, starting from the stock viral solution in the syringe, RSV undergoes three sequential processes: (i) aerosolization (t_0_), in which the liquid stock is atomized into droplets by the nebulizer under high-pressure compressed air; (ii) evaporation (t_1_), during which droplets dynamically shrink due to water loss as they equilibrate with the relative humidity (RH) of the dilution air; and (iii) early suspension (t_2_), in which bioaerosol particles reach equilibrium with the ambient RH and maintain stable physical properties. Sampling was conducted at the end of each stage to evaluate RSV survival.

A note is warranted regarding the selection of a 10 s evaporation period. The time required for droplets to evaporate and reach equilibrium with the environmental RH depends on initial size, chemical composition, and ambient RH. In this study, the sampling location was fixed to provide a uniform residence time of 10 s in the evaporation stage for all particles, independent of composition, initial size, and the two RH conditions examined (35% and 61%), to ensure experimental consistency. This duration was sufficient, as confirmed by SEM analysis of bioaerosol particles collected beyond the 10 s residence time, which showed no changes in phase or morphology (**Figure S1**). In addition, prior studies have reported that even larger droplets evaporate within <10 s under comparable RH conditions^25^, providing supporting evidence despite compositional differences.

#### Aerosol Particle Imaging

Aerosol particles were collected using an eight-stage Andersen Cascade Impactor (ACI) in a separate experimental setup that has been described in detail in our prior work ^26^ with detailed descriptions of the ACI configuration and SEM imaging procedures. In the present study, we incorporated a Nafion-tube humidifier coupled with a custom-built water bath to supply pre-conditioned dilution and central air, ensuring that the relative humidity within the ACI matched that of the flow-tube system. This configuration minimized additional evaporation within the impactor and enabled preservation of aerosol microenvironmental conditions during particle collection. Again, particle morphology and intraparticle chemical distributions were characterized using scanning electron microscopy (SEM) and energy-dispersive X-ray (EDX) line-scan analysis.

#### Virus preparation

RSV A2 purchased from ATCC (VR-1540) was propagated in HEp-2 cells (CCL-23) using Eagle’s minimum essential medium (EMEM; Corning, 10-009-CV) supplemented with 10% (v/v) heat-inactivated fetal bovine serum (FBS; Corning, 35-086-CV) and 1% Penicillin–Streptomycin (Corning, 30-002-CI). Cells were infected at a multiplicity of infection (MOI) of 0.05 plaque-forming units (PFU)/mL and incubated at 37 °C with 5% CO_2_.

When cytopathic effects (CPE) reached approximately 80–95% while the cells remained adherent (typically after 5–6 days), both the cell culture supernatant and attached cells were collected. The cells were scraped using a cell scraper, combined with the supernatant, and vortexed for 30 s to release an intracellular virus. The mixture was then centrifuged at 3000 × g for 10 min at 4 °C to remove the cell debris. The resulting solution is noted as virus medium below and used as base solution to form different stock solutions. The titer of the virus preparation was quantified by 50% Tissue Culture Infectious Dose (TCID_50_) as described below. We measured every batch for the experiment with the initial titer ranging from 3.16 × 10^6^ to 6.31 × 10^6^ TCID_50_/mL.

#### Solution preparation

Porcine stomach mucin (Millipore Sigma, M1778, Type III, bound sialic acid 0.5–1.5%, partially purified powder), and BSA (Fisher Scientific, BP9703100) were purchased and used without further modification. Three stock solutions were prepared: virus medium and virus medium supplemented with 3 g/L porcine mucin and 3 g/L of BSA separately. The chemical compositions of all solutions are reported in **Table S1-S4**.

The composition of the virus medium after virus propagation is expected to differ slightly from that of Eagle’s minimum essential medium (EMEM) supplemented with 10% (v/v) fetal bovine serum (FBS) and 1% Penicillin–Streptomycin. However, the chemical composition of the post-propagation media was not directly measured and was assumed to be similar to that of original virus propagation medium. Previous work by Niazi et al.^24^ reported that the chemical composition and concentrations of Opti-MEM supplemented with 2% FBS remained largely unchanged before and after RSV propagation. Based on this assumption, the chemical composition of the post-propagation virus medium was assumed to be the same as EMEM supplemented with 10% (v/v) FBS and 1% (v/v) Penicillin–Streptomycin. This assumption is further supported by the unaltered particle morphology and consistent energy-dispersive X-ray spectroscopy (EDX) spectra observed in SEM images (**Figure S2**).

#### TCID50 quantification

Infectious RSV titer was determined using a TCID_50_ assay. HEp-2 cells were cultured in Eagle’s minimum essential medium (EMEM) supplemented with 10% (v/v) heat-inactivated fetal bovine serum (FBS) and 1% (v/v) Penicillin–Streptomycin (Corning, 30-002-CI) in cell culture dishes (Falcon, 353003) and subsequently seeded into 96-well plates at ∼90% confluency.

Samples from both bulk solution and collection medium were serially diluted (10^−1^ to 10^−6^) in EMEM supplemented with 10%FBS and 1% Penicillin–Streptomycin. Each dilution was added to 8 replicate wells with a 100 µL per well. 96-wells Plates were incubated at 37 °C with 5% CO_2_ for 7 days. Following incubation, wells were examined for cytopathic effects (CPE), and the TCID_50_ values were calculated using the Reed–Muench method.

#### Digital Droplet PCR (ddPCR) Quantification

Viral RNA was extracted from aerosol samples collected in viral transport medium (VTM) using the QIAamp Viral RNA Mini Kit (Qiagen, Cat. No. 52904) following the manufacturer’s instructions and eluted in a final volume of 60 μL. Reverse transcription digital droplet PCR (RT-ddPCR) was performed using the One-Step RT-ddPCR Advanced Kit for Probes (Bio-Rad, Cat. No. 1864021) and ddPCR™ Expert Design Assays targeting RSV A and RSV B (Bio-Rad, Assay ID: dEXD77482599), according to the manufacturer’s protocol.

Each 22 μL reaction mixture contained 5 μL Supermix, 2 μL reverse transcriptase, 1 μL dithiothreitol, 1 μL assay mix, 5 μL RNA template, and 9 μL nuclease-free water. Droplets were generated using an automated droplet generator (Bio-Rad, Cat. No. 1864101). Thermal cycling was conducted under the following conditions: reverse transcription at 50°C for 60 min; enzyme activation at 95°C for 10 min; 40 cycles of denaturation at 94°C for 30 s and annealing/extension at 55°C for 1 min, with a ramp rate of 2°C/s; and enzyme deactivation at 98°C for 10 min, followed by a 4°C hold. Droplets were analyzed using a QX200™ Droplet Reader (Bio-Rad, Cat. No. 1864003), and absolute RNA copy numbers were calculated using QX Manager Software (version 2.3.1 standard edition, Bio-Rad). Additional details regarding reaction composition and assay performance are provided in **Table S10**.

#### Data analysis

Viral infectivity was quantified by normalizing infectious titer to viral RNA concentration to account for particle loss in the flow tube and potential losses during sampling and handling. This metric is referred to as the ratio of biological copies, *R*_*BC*_. Infectious virus was quantified by TCID_50_ (reported in TCID_50_ mL^−1^), and viral RNA concentration was measured by digital droplet PCR (reported in RNA copies μL^−1^).

To comparing RSV survival across aerosol stages, *R*_*BC*_ values were determined for bulk solutions and aerosol samples collected at residence times (*t*) of 0, 10, and 70 s, corresponding to the end of aerosolization, evaporation, and equilibrium stages, respectively. *R*_*BC*_ was calculated as:

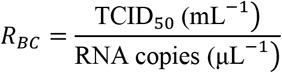

Relative survival ratios were defined by normalizing *R*_*BC,t*_, where *t* = 0, 10, and 70 s, to the corresponding bulk value (*R*_*BC,bulk*_):

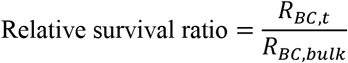

Stage-specific survival was further evaluated using ratios between stages. Survival during evaporation was quantified as *R*_*BC*,10 *s*_/*R*_*BC*,0 *s*_ for each given condition, and survival during the equilibrium stage was quantified as *R*_*BC*,70 *s*_/*R*_*BC*,10 *s*_.

#### Statistical analysis

Statistical analyses were performed to compare survival ratios among groups under each relative humidity condition. For each relative humidity level, data were analyzed separately using one-way analysis of variance (one-way ANOVA), followed by Tukey’s multiple-comparison post hoc test to identify significant differences between groups. Comparisons were performed separately for the 35% RH and 61% RH conditions. Statistical significance was defined as p < 0.05.

## Results

RSV infectivity in the flow tube depends strongly on residence time, chemical composition, and relative humidity (RH). Relative survival ratios for three residence times (0, 10 s, and 70 s), two RH conditions (35% and 61%), and three solution compositions (virus medium only, and virus medium supplemented with 3 g L^−1^ mucin or 3 g L^−1^ BSA) are shown in **Figure 2**.

**Figure 2.**
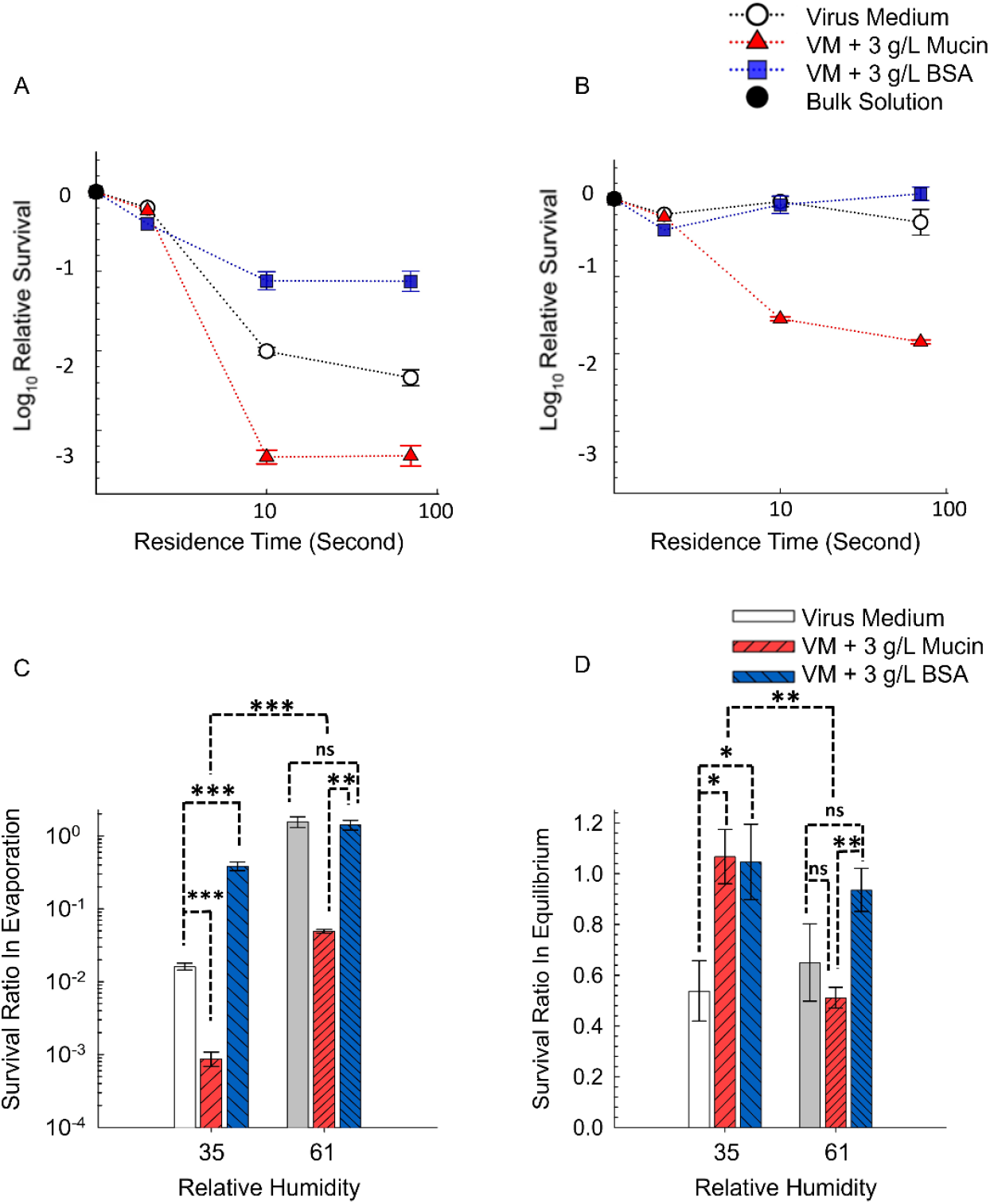
RSV relative survival ratio at different suspension times in the flow tube at 35% and 61% relative humidity for aerosols generated from solutions with varying chemical compositions: virus medium alone and virus medium supplemented with 3 g L^−1^ mucin and 3 g L^−1^ BSA separately. Survival normalized to the biological infectivity of the bulk solution at 35% RH (A) and at 61% (B); Survival ratio during evaporation, with measurements at 10 s normalized to the aerosolization stage (C); Survival ratio at equilibrium, with measurements at 70 s normalized to the 10 s residence time (D).

### RSV rapidly loses infectivity during evaporation at low RH

As shown in Figure 2A, RSV experiences limited loss during aerosolization, with an average reduction of approximately 50%, indicating that the custom-built large aerosol generator largely preserves viral infectivity during this stage. In contrast, substantial loss occurs during the evaporation stage at low RH. Across all three media, relative survival ratios decrease by one to three orders of magnitude. Among these, BSA-supplemented particles exhibit the highest survival, while mucin-supplemented particles show the lowest survival, approximately two orders of magnitude lower than BSA-supplemented particles.

### RSV survival is enhanced at 61% RH with compositional dependence

At intermediate RH (∼61%) (Fig. 2B), RSV survival at both 10 s and 70 s is significantly higher than at 35% RH for all media, with improvements of approximately one order of magnitude. For virus medium and BSA-supplemented particles, RSV experiences minimal loss, with infectivity remaining close to that at the aerosolization stage. In contrast, mucin-supplemented particles consistently exhibit lower survival than the other two media during both evaporation and equilibrium stages.

### RSV survival during evaporation strongly depends on chemical composition

As shown in Figure 2C, at low RH, RSV survival follows the order: BSA-supplemented > virus medium > mucin-supplemented particles, with all differences being statistically significant. At intermediate RH, all media exhibit higher survival than at low RH, and virus medium and BSA-supplemented particles again show higher survival than mucin-supplemented particles.

### RSV infectivity loss during equilibrium is smaller compared to the evaporation stage

As shown in **Figure 2D**, at equilibrium (70 s), after evaporation is complete and particles reach stable physicochemical states, RSV infectivity remains largely unchanged, with relative survival ratios approaching unity. This contrast with the substantial losses observed during evaporation indicates that viral inactivation is dominated by early-stage processes. However, two notable differences are observed. First, at 35% RH, virus medium particles exhibit lower survival than both mucin- and BSA-supplemented particles, indicating a protective effect of added organics under dry conditions. Second, at 61% RH, mucin-supplemented particles exhibit slightly lower survival than virus medium and BSA-supplemented particles, suggesting that mucin influences RSV survival differently from BSA at this RH.

### SEM images

The observed variations in RSV survival across stages (evaporation vs. equilibrium), RH conditions (35% vs. 61%), and chemical compositions (virus medium particles versus mucin- and BSA-supplemented particles) likely arise from complex microenvironmental conditions within the virus-laden particles. To probe these conditions, bioaerosol particles were collected at a residence time of 10 s under controlled RH and subsequently transported to the imaging facility for SEM analysis, during which the RH decreased to ambient levels (∼30–40%). As shown in **Figure 3**, particle morphologies are diverse and strongly dependent on both chemical composition and RH.

**Figure 3.**
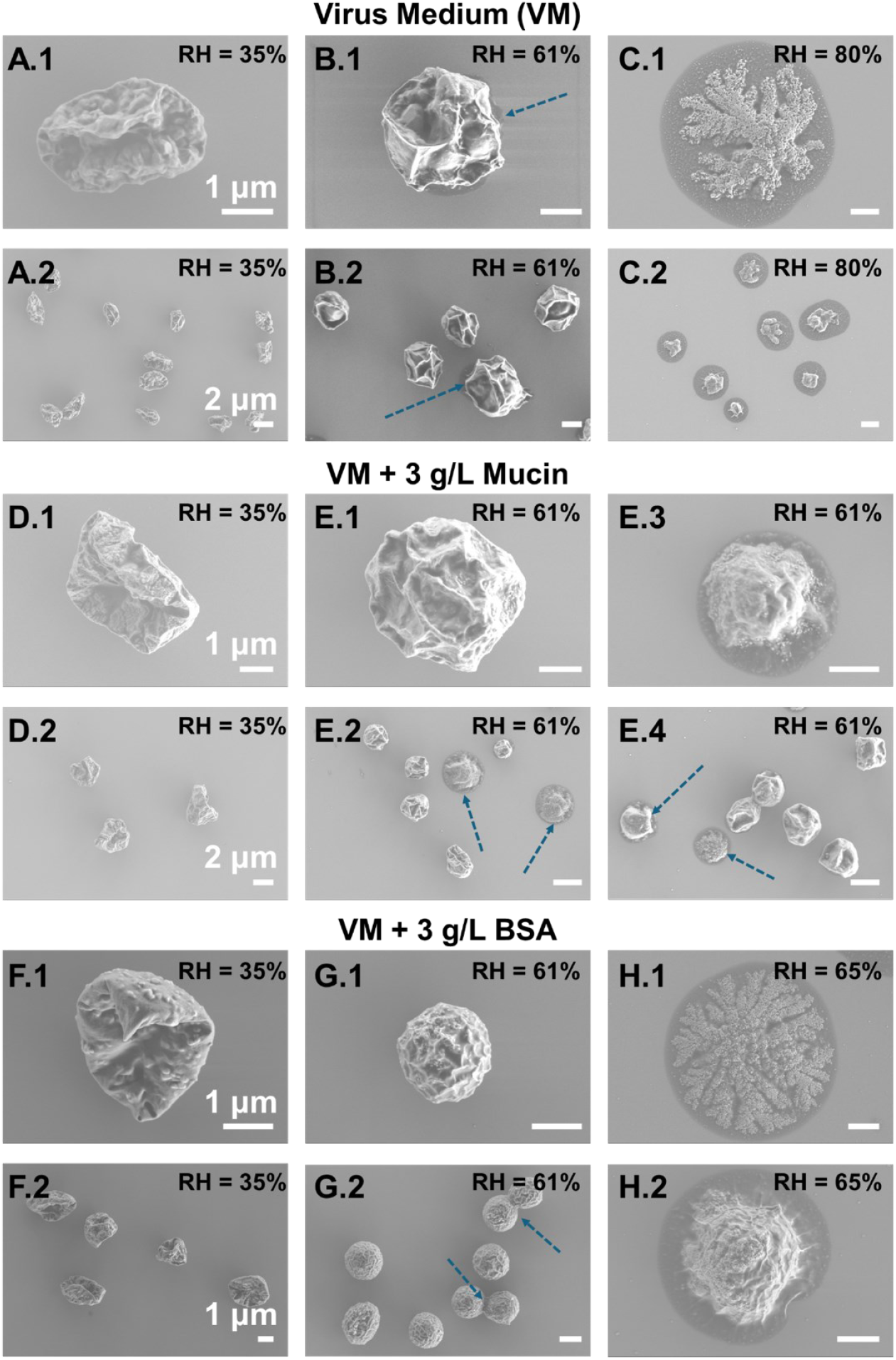
Bioaerosol particle morphology depends strongly on both chemical composition and relative humidity (RH). SEM images of: (A) virus medium (VM) particles at 35% RH (A1, A2), 61% RH (B1, B2), and 80% RH (C1, C2); (B) mucin-supplemented particles at 35% RH (D1, D2) and 61% RH (E1–E4); (C) BSA-supplemented particles at 35% RH (F1, F2), 61% RH (G1, G2), and 65% RH (H1, H2).

### Efflorescence at Low RH Is Associated with Reduced RSV Infectivity

At low RH (35%), bioaerosol particles generated from virus medium, as well as mucin and BSA-supplemented media, likely undergo efflorescence. First, the base medium (EMEM with 10% FBS and 1% Penicillin) contains a significant inorganic fraction, with NaCl as the dominant salt (∼6.8 g L^−1^), which is expected to crystallize under low RH conditions. Second, the high evaporation rates at 35% RH produce particles with irregular morphologies that deviate from spherical shapes, exhibiting features such as cavities, dents, and folds. These structures are consistent with surface restructuring associated with efflorescence during rapid drying. Third, crystalline structures are directly observed in SEM images. In VM particles, coarse crystals are embedded within a thin organic-rich layer (**Figure 3A1**), whereas in BSA-supplemented particles, finer crystals are observed and appear more exposed at the particle surface (**Figure 3F1**). Together, these morphological and compositional observations support the occurrence of efflorescence under low RH conditions.

Consistent with these observations, RSV infectivity is markedly reduced at 35% RH. This reduction is likely linked to efflorescence-induced changes in particle physicochemical properties. Supporting this interpretation, Oswin et al.^15^ demonstrated that salt crystallization during efflorescence can significantly enhance viral inactivation of SARS-CoV-2 by cycling RH between ∼70% and 40%. While the exact mechanisms may differ between systems, our results suggest that efflorescence is an important contributing factor to the reduced RSV survival observed under low RH condition.

### Slower Evaporation and Higher Water Content at Intermediate RH Are Associated with Higher RSV Infectivity

During the evaporation stage, RSV survival is higher at intermediate RH (∼61%) than at low RH (35%) for all three media tested: virus medium alone and virus medium supplemented with 3 g L^−1^ BSA or mucin (**Figure 2C**). These differences are statistically significant for each medium. Enhanced survival is likely associated with slower evaporation rates and the presence of a viscous or semi-solid particle phase with higher water content at intermediate RH.

As shown in **Figure 3**, SEM observations indicate that particles become more spherical at intermediate RH. In particular, BSA-supplemented particles exhibit highly spherical morphologies, suggesting that slower evaporation allows nonvolatile ions and molecules sufficient time to redistribute more homogeneously and form stable three-dimensional structures.

Particles at intermediate RH are also consistent with a viscous or semi-solid state with higher water content. For VM particles, the organic-rich phase appears capable of flowing, resulting in circular residue patterns upon deposition (**Figure 3B1**). Mucin-supplemented particles show a noticeable fraction of spreading particles (**Figure 3E2–3E4**) at intermediate RH, a behavior not observed at low RH, further indicating increased water content. Similarly, BSA-supplemented particles exhibit occasional bridging between neighboring particles (**Figure 3G2**), suggesting a viscous, hydrated state that allows particle–particle coalescence.

To further distinguish particle phase behavior from potential artifacts during deposition or post-collection drying, additional experiments were conducted at higher RH. Specifically, RH was increased to 80% for virus medium particles and to 65% for BSA-supplemented particles. Under these conditions, particles exhibited larger projected areas and pronounced spreading upon deposition, consistent with fully liquid droplets. After evaporation during sample handling, these droplets formed characteristic circular residue patterns and reorganized into structures such as dendritic (“tree-like”) or central “mountain-like” features ^27^. All these features are very different from particles at 61% RH and confirms that particles are at viscous/semi-solid in the aerosol state with relatively high water content.

### Lower RSV Survival at Equilibrium is Associated with Core–Shell Inorganic–Organic Separation in VM Particles at 35% RH

Compared to the relatively uniform elemental distributions observed in organic-supplemented systems (**Figure 3**), VM particles exhibit clear inorganic–organic separation and a distinct core– shell structure at both RH conditions investigated (**Figure 4**). At low RH, fine crystalline structures are observed and appear to be coated by additional components, likely organics (**Figure 4A1**). This interpretation is supported by several observations: (1) differences in contrast between bare crystals and crystals covered by a thin organic layer; (2) spatial separation between inorganic (Na, Cl) and organic (C, O) elemental peaks in EDX maps (**Figure 4A2**); and (3) relatively uniform C and O signal intensities across the particle (**Figure 4B2**), consistent with an organic coating over inorganic cores. In contrast, if the components were well mixed, elemental signals would show co-located peaks due to cumulative contributions along the projection direction.

**Figure 4.**
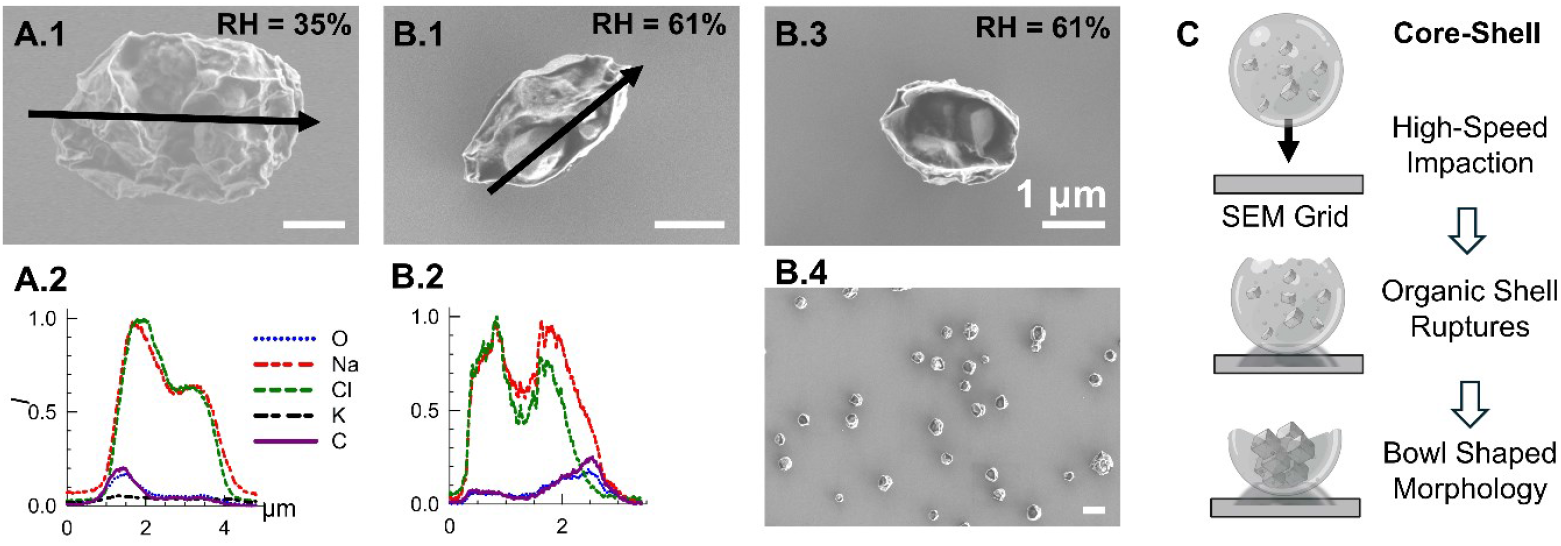
Core–shell structures with an inorganic-rich core and an organic-rich shell in particles generated from virus medium. (A) Particle morphology and elemental distribution at 35% RH; (B) Bowl-shaped morphology at 61% RH, with visible large crystals enclosed within the organic shell; (C) Schematic illustration of the formation of the bowl-shaped structure at 61% RH.

Consistent with this structural difference, RSV inactivates more rapidly in VM particles than in organic-supplemented particles at equilibrium (**Figure 2D**). This difference may be associated with the localization of organic material near the particle surface, assuming virus colocalized with organic contents, which may place viruses closer to the particle–air interface. Coupled with efflorescence in the particle core and reduced local water retention after drying, this configuration may increase exposure to interfacial stresses and contribute to enhanced viral inactivation. In contrast, mucin and BSA-supplemented particles exhibit more homogeneous chemical distributions, suggesting that viruses may be more uniformly distributed within the particle and not preferentially located near the surface.

### Inorganic-organic separation does not suppress RSV survival at 61% RH

At intermediate RH (61%), the core–shell structure of VM particles becomes more evident. Due to the higher water content at this RH, the organic layer is viscous and capable of flowing. Upon impaction, the viscous shell may rupture, exposing the inorganic core to the environment, where it subsequently completes crystallization (**Figure 4B3**). Whether the inorganic core exists as a fully liquid phase or contains a few nucleation sites (inclusions ^28^) in the aerosol state remains unclear. It is possible that such inclusions are already present in the aerosol state and evolve into well-defined crystals during subsequent drying on the substrate. This process results in the observed bowl-shaped particle morphology via SEM imaging. Although not all particles experience sufficient impaction energy to rupture the shell, the significant fraction of ruptured particles supports the presence of inorganic–organic phase separation with a core–shell configuration.

Despite this core-shell structure, RSV survival at intermediate RH remains high in VM particles, with survival comparable to that observed in BSA-supplemented particles in both the evaporation and equilibrium stages. This behavior is likely associated with the higher water content at this RH; although inorganic–organic phase separation is present, the elevated water content appears to mitigate its impact, allowing the virus to remain relatively stable.

## Discussions

### BSA Preserves RSV at Both RH Conditions

BSA seems to preserve RSV well at both RH values and in both evaporation and equilibrium stages. Similar protection has been reported for the surrogate virus bacteriophage ϕ6, where BSA improves infectivity in both aerosol^29^ and droplet-on-surface^21^ conditions. In this study, at low RH, although efflorescence occurs in both VM and BSA-supplemented particles, the presence of BSA is associated with a more homogeneous intraparticle chemical distribution. This may allow viruses to be more evenly distributed within the particle, potentially reducing their localization at the particle–air interface, where interfacial stresses are expected to be more pronounced. At intermediate RH, homogenous intraparticle chemical distributions might be less important compared to the particle water content. Nevertheless, the exact mechanisms by which BSA interacts with viruses under these dynamically evolving microenvironmental conditions remain uncertain, and further study is needed to fully elucidate these processes.

### RSV Infectivity Is Lower in Mucin-Supplemented Particles at Both RH Conditions

Although mucin contains a high protein content, it behaves differently from BSA. In particular, RSV survival in mucin-supplemented particles shows a reduction of one to two orders of magnitude compared to BSA-supplemented particles during the evaporation stage.

Interestingly, once particles reach a dry and stable equilibrium state at low RH, RSV survival becomes similar between mucin- and BSA-supplemented particles. This observation is consistent with SEM and elemental analyses indicating that both particle types undergo efflorescence and exhibit relatively well-mixed intraparticle chemical distributions under these conditions. At intermediate RH (61%), mucin-supplemented particles also show a hydrated and morphologically similar state compared to the other particle types; however, differences in RSV survival persist.

Taken together, these observations suggest that mucin may reduce RSV survival in the presence of water; however, the underlying mechanism remains unclear. Previous work by Wardzala et al.^30^ has shown that mucin can interact with viral surface proteins through glycan-mediated binding, potentially inhibiting viral infection. To date, similar studies specifically examining interactions between mucin and RSV are limited, and the chemical and biological interactions between mucin and RSV therefore remain poorly understood^31^. Notably, mucin has been reported to enhance viral survival in some systems, including influenza A virus in droplet-on-surface conditions^20^ and coronaviruses^28^ and bacteriophage ϕ6^29^ in aerosol systems. In contrast, other studies have shown that mucin can reduce viral survival in aerosols compared to FBS-supplemented media^32^. Taken together, these findings indicate that the effects of mucin on viral survival are complex and system dependent. Further investigation is therefore needed to clarify the role of mucin in RSV survival under aerosol conditions.

### Significance for short-range transmission via coughing

In this study, we employed a flow-tube system that enables precise control of short suspension times by adjusting the carrier-gas flow rate^33^. The total suspension time was limited to 70 s, yet allow simultaneous sampling with two BioSamplers for characterization of the equilibrium stage. This duration was sufficient to clearly separate the evaporation stage from the equilibrium stage, facilitating mechanistic understanding of virus survival in bioaerosol particles.

Under a cough-generated respiratory puff, aerosols can travel beyond approximately 2 m within 10 s and up to ∼3 m within 60 s^34^, making the timescales investigated here directly relevant to coughing and sneezing events^8^. Distinguishing between evaporation-stage and suspension-stage inactivation is therefore critical for understanding RSV transmission. Transmission among vulnerable populations, such as infants^35^, transplant patients^36,37^ and elderly individuals in nursing homes ^38–42^, is likely dominated by short-range exposure, where respiratory bioaerosols are either still evaporating or have just reached equilibrium prior to inhalation by a susceptible individual.

### Limitations

This study has several limitations that should be considered when interpreting the results. First, bioaerosol sampling was conducted at only three discrete residence times (0, 10, and 70 s), which enabled separation of the aerosolization, evaporation, and equilibrium stages but provided limited temporal resolution within each stage. As a result, rapid and dynamic changes in viral survival during the evaporation process may not be fully resolved.

Second, the experimental design focuses on short suspension times to capture evaporation-driven processes, and long-term aerosol suspension was not investigated. Therefore, the findings are most directly applicable to short-range transmission scenarios and may not extend to far-field or long-range transport, where additional physicochemical processes may influence viral survival.

Third, the interpretation of mucin effects remains challenging due to the coupled influence of mucin and inorganic salts on particle physicochemical properties. Interactions among mucin concentration, salt content, and phase behavior complicate the separation of biological effects from physical processes. Further studies that systematically vary mucin and salt concentrations are needed to better isolate their individual and combined roles in influencing viral survival.

Fourth, the chemical compositions used in this study, including virus medium and organic-supplemented formulations, represent simplified surrogates of respiratory fluids. Real respiratory aerosols, such as saliva and airway lining fluid, are more chemically complex and variable in composition. Therefore, caution is warranted when extrapolating these findings to real-world conditions, and additional work using more representative biological fluids is needed to fully evaluate RSV survival in authentic respiratory microenvironments.

## Conclusions

In this study, we investigated RSV survival in bioaerosols across aerosolization, evaporation, and early suspension stages under controlled relative humidity and chemical compositions. Our results show that RSV inactivation is strongly stage dependent, with the majority of infectivity loss occurring during the evaporation stage. At low RH (35%), rapid evaporation and efflorescence lead to substantial inactivation, whereas at intermediate RH (61%), slower evaporation and higher water content result in significantly enhanced survival.

Particle phase state plays a central role in mediating these outcomes. SEM and elemental analyses indicate that low RH conditions favor crystalline, effloresced particles, while intermediate RH produces viscous or semi-solid particles with higher water content. These differences in microenvironmental conditions are closely linked to viral survival. In addition, inorganic–organic phase separation in virus medium particles forms a core–shell structure that is associated with reduced survival at low RH, likely due to increased interfacial exposure and reduced local water retention. In contrast, this structural effect is diminished at intermediate RH, where elevated water content dominates.

Chemical composition further modulates RSV survival. BSA consistently enhances viral stability across RH conditions, potentially through improved water retention and more homogeneous intraparticle distributions. In contrast, mucin exhibits a more complex effect, reducing RSV infectivity in hydrated states despite similar particle morphologies. This suggests that additional studies exploring a range of mucin concentrations, as well as potential biological and molecular interactions beyond purely physical mechanisms, are needed.

Overall, these results demonstrate that RSV survival in aerosols is governed by the interplay between evaporation dynamics, phase transitions, and particle composition. Importantly, the dominant role of early evaporation-stage processes indicates that viral viability is largely determined within the first seconds of airborne transport. These findings provide mechanistic insight into RSV stability under conditions relevant to short-range transmission and emphasize the importance of aerosol microenvironmental evolution in controlling viral infectivity.

## Acknowledgments

The authors acknowledge the Clean Room and Imaging Core Facility at the University of Texas at Dallas for providing access to SEM instrumentation.

