## Supplemental Information for "Intermediate Relative Humidity Preserves Respiratory Syncytial Virus via a Semi-Solid Bioaerosol State"

**Short Title: Semi-Solid Bioaerosols Enhance RSV Survival at Intermediate RH**

Yuhui Guo^a^†, Deepak Sapkota^a^†, Ajay Sajan^a^, HoangDinh Huynh^b^, Imtiaz Taimoor^a^, Jeffrey Kahn ^b,c^, Hui Ouyang^a^*

^a^ Department of Mechanical Engineering, University of Texas at Dallas, Richardson, TX, USA

^b^ Department of Pediatrics, University of Texas Southwestern Medical Center, Dallas, TX, USA

^c^ Department of Microbiology, University of Texas Southwestern Medical Center, Dallas, TX, USA

† These authors contributed equally.

*Corresponding author: Hui Ouyang

**Table S1: Chemical Composition of Eagle’s Minimum Essential Medium (EMEM) (Corning, NY, USA, Cat. No: 10-009-CV).**

| **Chemical Compound** | **Concentration (g/L)** |
| --- | --- |
| CaCl_2_ (anhydrous) | 0.2 |
| KCl | 0.4 |
| MgSO_4_ (anhydrous) | 0.0977 |
| NaCl | 6.8 |
| NaH_2_PO_4_.H_2_O | 0.14 |
| NaHCO_3_ | 1.5 |
| L-Alanine | 0.0089 |
| L-Asparagine.H_2_O | 0.015 |
| L-Aspartic acid | 0.0133 |
| L-Arginine.HCl | 0.1264 |
| L-Cystine .2HCl | 0.0312 |
| L-Glutamic acid | 0.0147 |
| L-Glutamine | 0.292 |
| Glycine | 0.0075 |
| L-Histidine.HCl.H_2_O | 0.0419 |
| L-Isoleucine | 0.0525 |
| L-Leucine | 0.0525 |
| L-Lysine. HCl | 0.0725 |
| L-Methionine | 0.015 |
| L-Phenylalanine | 0.0325 |
| L-Proline | 0.0115 |
| L-Serine | 0.0105 |
| L-Threonine | 0.0476 |
| L-Tryptophan | 0.01 |
| L-Tyrosine.2Na.2H_2_O | 0.0519 |
| L-Valine | 0.0468 |
| D-Calcium pantothenate | 0.001 |
| Choline chloride | 0.001 |
| Folic acid | 0.001 |
| i-Inositol | 0.002 |
| Nicotinamide | 0.001 |
| Pyridoxine. HCl | 0.001 |
| Riboflavin | 0.0001 |
| Thiamine.HCl | 0.001 |
| D-Glucose | 1 |
| Phenol red.Na | 0.01 |
| Sodium Pyruvate | 0.110 |

**Table S2: Chemical Composition Cell Culture Medium**

| **Chemical Compound** | **Concentration (v/v)** |
| --- | --- |
| Eagle’s Minimum Essential Medium (EMEM) | 89% |
| FBS (Corning, NY, USA Cat. No: 35-086-CV) | 10 % |
| Penicillin–Streptomycin (Corning, 30-002-CI) | 1% |

**Table S3: Chemical Composition of Virus Medium + 3g/L Mucin**

| **Chemical Compound** | **Concentration (g/L)** |
| --- | --- |
| Virus Medium | RSV Propagated Cell Culture Medium |
| Porcine Stomach Mucin (Millipore Sigma, M1778, Type III, bound sialic acid 0.5–1.5%, partially purified powder) | 3 |

**Table S4: Chemical Composition of Virus Medium + 3g/L BSA**

| **Chemical Compound** | **Concentration (g/L)** |
| --- | --- |
| Virus Medium | RSV Propagated Cell Culture Medium |
| Bovine Serum Albumin (Fisher Scientific, BP9703100) | 3 |

**Table S5: The Concentration of extracted RNA and TCID50/mL in the bulk solution**

| **Media** | **Bulk Solution** | |
| --- | --- | --- |
|  | **RNA(Copies/μL)** | **TCID50/mL** |
| Virus Media | 5.11E+04 | 5.31E+06 |
| Virus Media + 3 g/L Mucin | 5.02E+04 | 3.16E+06 |
| Virus Media + 3 g/L BSA | 4.79E+04 | 4.64E+06 |

**Table S6: The Concentration of extracted RNA and TCID50/mL at *t* = 0 s (Aerosolization)**

| **Relative Humidity** | **T0** | |
| --- | --- | --- |
|  | **RNA(Copies/μL)** | **TCID50/mL** |
| 91% | 7.25E+03 | 4.53E+05 |
|  | 6.09E+03 | 7.57E+04 |
|  | 5.52E+03 | 2.17E+05 |

**Table S7: The Concentration of extracted RNA and TCID50/mL at the *t* = 10 s (Completion of Evaporation)**

| **Media** | **Relative Humidity** | **T10** | |
| --- | --- | --- | --- |
|  |  | **RNA(Copies/μL)** | **TCID50/mL** |
| Virus Media | 35% | 3.27E+03 | 3.22E+03 |
| Virus Media + 3 g/L Mucin |  | 7.23E+03 | 1.39E+03 |
| Virus Media + 3 g/L BSA |  | 6.03E+03 | 9.34E+04 |
| Virus Media | 61% | 2.82E+03 | 2.68E+05 |
| Virus Media + 3 g/L Mucin |  | 7.42E+03 | 2.04E+04 |
| Virus Media + 3 g/L BSA |  | 4.81E+03 | 3.16E+05 |

**Table S8: The Concentration of extracted RNA and TCID50/mL at *t* = 70 s (Equilibrium)**

| **Media** | **Relative Humidity** | ***t* = 70 s** | |
| --- | --- | --- | --- |
|  |  | **RNA(Copies/μL)** | **TCID50/mL** |
| Virus Media | 35% | 2.01E+03 | 1.05E+03 |
| Virus Media + 3 g/L Mucin |  | 3.26E+03 | 6.06E+02 |
| Virus Media + 3 g/L BSA |  | 3.23E+03 | 5.33E+04 |
| Virus Media | 61% | 6.26E+02 | 3.91E+04 |
| Virus Media + 3 g/L Mucin |  | 3.80E+03 | 5.02E+03 |
| Virus Media + 3 g/L BSA |  | 3.57E+03 | 2.20E+05 |

**Table S9. One-way ANOVA with Tukey’s multiple-comparison analysis of log10-transformed relative survival ratios under two RH conditions and two aerosol stages.**

Statistical analyses were performed on log10-transformed relative survival ratios. Mean differences are reported as absolute mean differences. Comparisons among media conditions within the same RH condition and aerosol stages were performed using one-way ANOVA followed by Tukey’s multiple-comparison test. Comparisons between 35% RH and 61% RH for the same medium and aerosol stage were performed using two-tailed t-tests. Significance levels are indicated as ns, not significant; *P < 0.05; **P < 0.01; ***P < 0.001. VM is virus medium.

| **RH condition** | **Aerosol stage** | **Comparison** | **Mean**  **Difference** | **Significant?** | **Summary** | **P Value** |
| --- | --- | --- | --- | --- | --- | --- |
| 35% RH | Evaporation | 35% VM vs 35% Mucin | 0.0154 | Yes | *** | <0.001 |
| 35% RH | Evaporation | 35% BSA vs 35% VM | 0.37 | Yes | *** | <0.001 |
| 35% RH | Evaporation | 35% BSA vs 35% Mucin | 0.386 | Yes | *** | <0.001 |
| 61% RH | Evaporation | 61% VM vs 61% Mucin | 1.518 | Yes | ** | 0.002 |
| 61% RH | Evaporation | 61% VM vs 61% BSA | 0.147 | No | ns | 0.896 |
| 61% RH | Evaporation | 61% BSA vs 61% Mucin | 1.371 | Yes | ** | 0.004 |
| 35% vs 61% RH | Evaporation | 61% Mucin vs 35% Mucin | 0.0485 | Yes | *** | <0.001 |
| 35% RH | Equilibrium | 35% Mucin vs 35% VM | 0.529 | Yes | * | 0.014 |
| 35% RH | Equilibrium | 35% BSA vs 35% VM | 0.508 | Yes | * | 0.032 |
| 35% RH | Equilibrium | 35% Mucin vs 35% BSA | 0.021 | No | ns | 0.925 |
| 61% RH | Equilibrium | 61% VM vs 61% Mucin | 0.138 | No | ns | 0.412 |
| 61% RH | Equilibrium | 61% BSA vs 61% VM | 0.287 | No | ns | 0.155 |
| 61% RH | Equilibrium | 61% BSA vs 61% Mucin | 0.425 | Yes | ** | 0.005 |
| 35% vs 61% RH | Equilibrium | 35% Mucin vs 61% Mucin | 0.556 | Yes | ** | 0.004 |

**Table S10: Reverse Transcription digital droplet PCR Reaction Setup and Thermal Cycling Conditions**

| **Component** | **Volume per reaction (µL)** | **Final concentration** |
| --- | --- | --- |
| Supermix | 5 | 1× |
| Reverse transcriptase | 2 | 20 U/µL |
| 300 mM DTT | 1 | 15 mM |
| 20× ddPCR assay | 1 | 1× |
| RNA sample | 5 | Variable |
| RNase/DNase-free water | 7 | - |
| **Total volume** | **20** | - |

| **Cycling step** | **Temperature (°C)** | **Time** | **Ramp rate** | **Number of cycles** |
| --- | --- | --- | --- | --- |
| Reverse transcription | 50 | 60 min | 2 °C/s | 1 |
| Enzyme activation | 95 | 10 min | 2 °C/s | 1 |
| Denaturation | 94 | 30 s | 2 °C/s | 40 |
| Annealing/extension | 55 | 1 min | 2 °C/s | 40 |
| Enzyme deactivation | 98 | 10 min | 2 °C/s | 1 |
| Hold | 4 | Infinite | 1 °C/s | 1 |

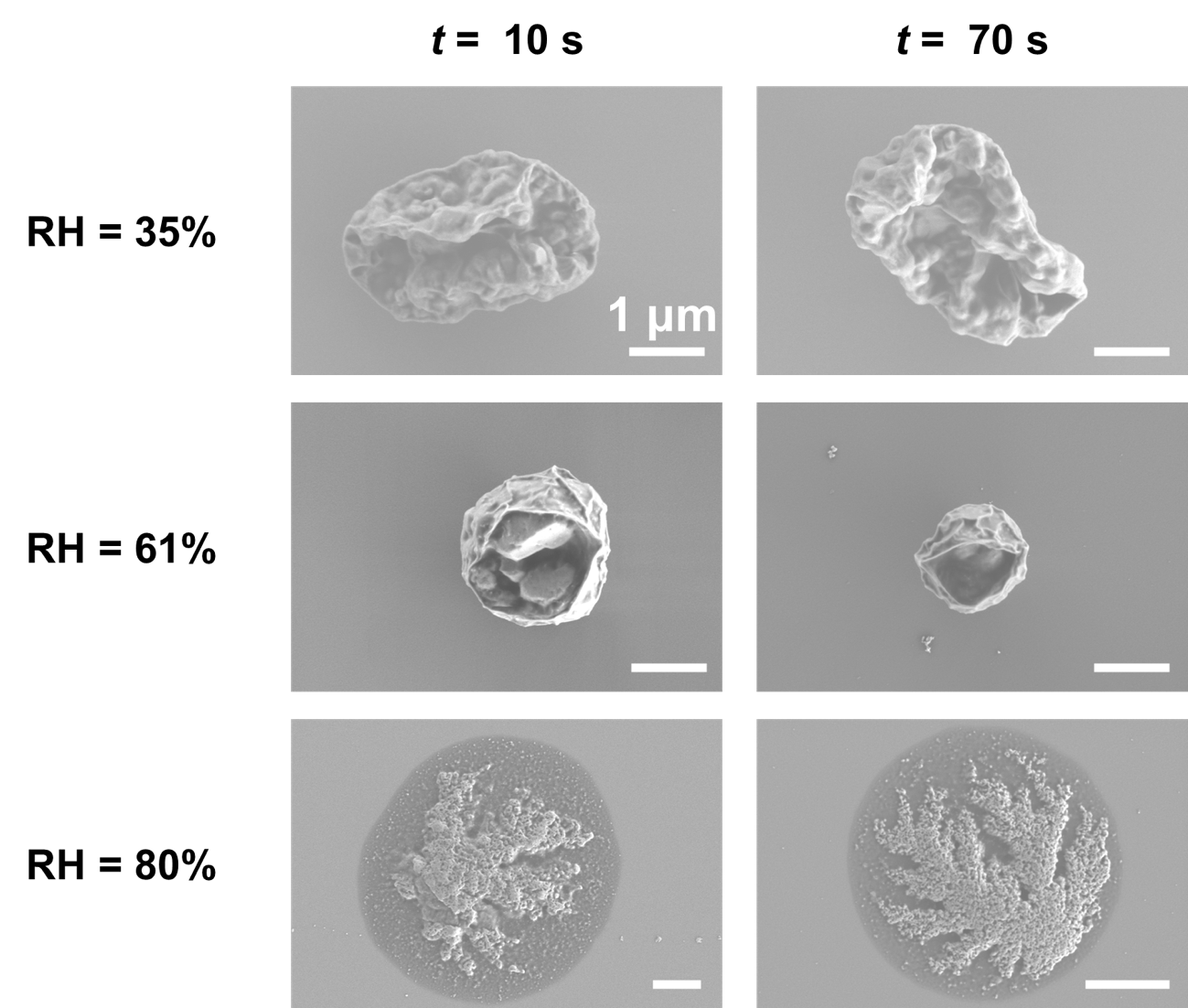

**Figure S1**. Virus Medium (VM) aerosols at 10 s and 70 s residence time for different relative humidities

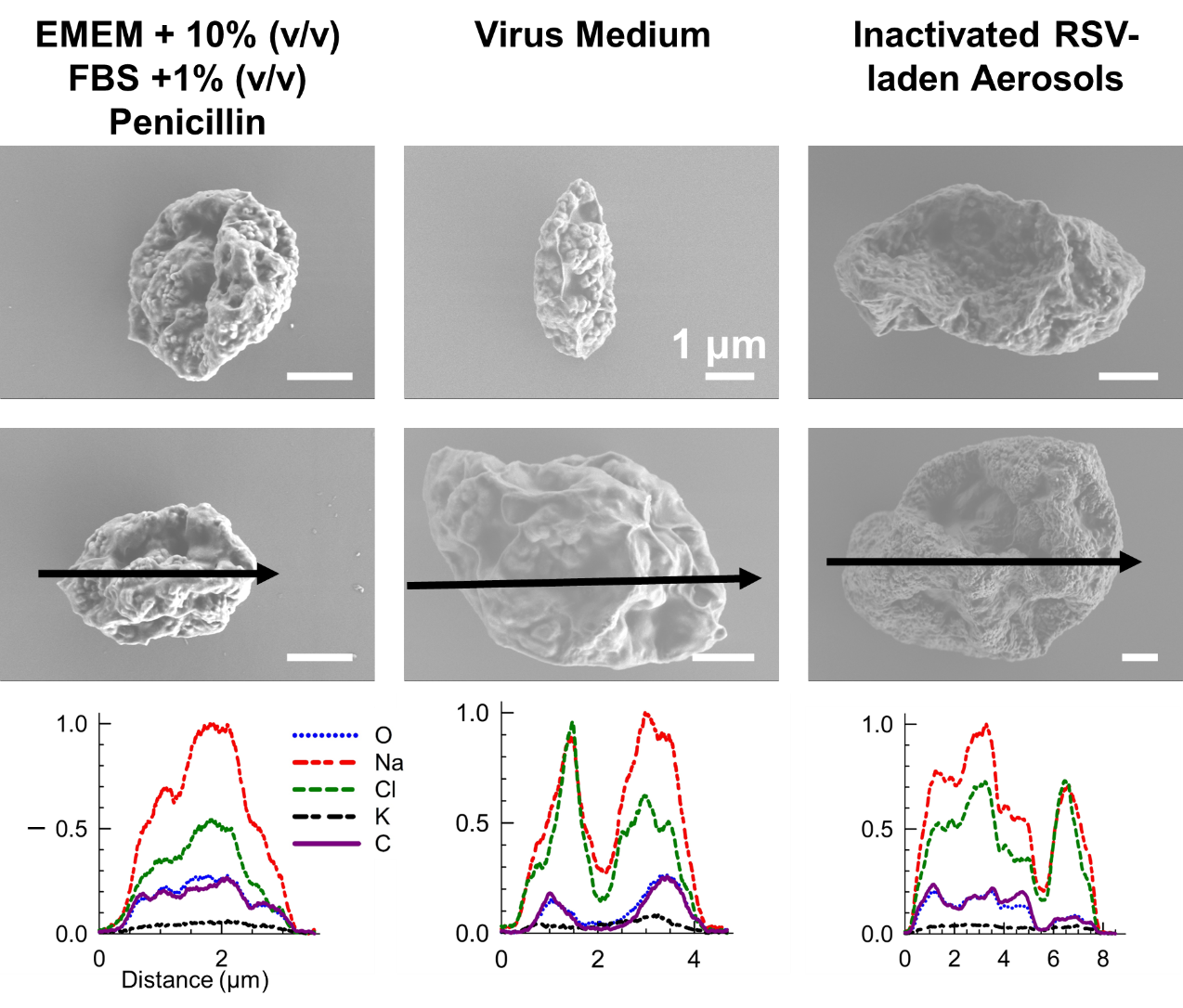

**Figure S2**. Morphology and chemical distribution of EMEM supplemented with 10% (v/v) FBS and 1% (v/v) Penicillin, Virus Medium and Heat inactivated Virus Medium with RSV.
